# Immunological correlates of protection mediated by a whole organism *Cryptococcus neoformans* vaccine deficient in chitosan

**DOI:** 10.1101/2024.06.12.598760

**Authors:** Charles A. Specht, Ruiying Wang, Lorena V. N. Oliveira, Maureen M. Hester, Christina Gomez, Zhongming Mou, Diana Carlson, Chrono K. Lee, Camaron R. Hole, Woei C. Lam, Rajendra Upadhya, Jennifer K. Lodge, Stuart M. Levitz

## Abstract

The global burden of infections due to the pathogenic fungus *Cryptococcus* is substantial in persons with low CD4^+^ T cell counts. Previously, we deleted three chitin deacetylase genes from *C. neoformans* to create a chitosan-deficient, avirulent strain, designated *cda1Δ2Δ3Δ* which, when used as a vaccine, protected mice from challenge with virulent *C. neoformans* strain KN99. Here, we explored the immunological basis for protection. Vaccine-mediated protection was maintained in mice lacking B cells or CD8^+^ T cells. In contrast, protection was lost in mice lacking α/β T cells or CD4^+^ T cells. Moreover, CD4^+^ T cells from vaccinated mice conferred protection upon adoptive transfer to naive mice. Importantly, while monoclonal antibody-mediated depletion of CD4^+^ T cells just prior to vaccination resulted in complete loss of protection, significant protection was retained in mice depleted of CD4^+^ T cells after vaccination, but prior to challenge. Vaccine-mediated protection was lost in mice genetically deficient in IFNγ, TNFα, or IL-23p19. A robust influx of leukocytes and IFNγ- and TNFα-expressing CD4^+^ T cells was seen in the lungs of vaccinated and challenged mice. Finally, a higher level of IFNγ production by lung cells stimulated ex vivo correlated with lower fungal burden in the lungs. Thus, while B cells and CD8^+^ T cells are dispensable, IFNγ and CD4^+^ T cells have overlapping roles in generating protective immunity prior to *cda1Δ2Δ3Δ* vaccination. However, once vaccinated, protection becomes less dependent on CD4^+^ T cells, suggesting a strategy for vaccinating HIV^+^ persons prior to loss of CD4^+^ T cells.

**Importance:** The fungus *Cryptococcus neoformans* is responsible for >100,000 deaths annually, mostly in persons with impaired CD4^+^ T cell function such as AIDS. There are no approved human vaccines. We previously created a genetically engineered avirulent strain of *C. neoformans*, designated *cda1Δ2Δ3Δ*. When used as a vaccine, *cda1Δ2Δ3Δ* protects mice against a subsequent challenge with a virulent *C. neoformans* strain. Here, we defined components of the immune system responsible for vaccine-mediated protection. We found that while B cells and CD8^+^ T cells were dispensible, protection was lost in mice genetically deficient in CD4^+^ T cells, and the cytokines IFNγ, TNFα, or IL-23. A robust influx of cytokine-producing CD4^+^ T cells was seen in the lungs of vaccinated mice following infection. Importantly, protection was retained in mice depleted of CD4^+^ T cells following vaccination, suggesting a strategy to protect persons who are at risk for future CD4^+^ T cell dysfunction.

## Introduction

Cryptococcosis, due to the encapsulated species of fungi including *C. neoformans* and *C. gattii*, is a major cause of morbidity and mortality worldwide. Human exposure is thought to mainly occur following inhalation of airborne organisms. A pulmonary infection may result which is often asymptomatic. In the absence of effective host defenses, infection can spread locally and also disseminate, most often to the central nervous system where it causes meningoencephalitis. Most persons with cryptococcosis have quantitative or qualitative CD4^+^ T cell dysfunction. The estimated global burden of cryptococcosis is 194,000 incident cases per year with 147,000 deaths (1). An estimated 19% of AIDS-related deaths are due to cryptococcal meningitis (2). Other immunosuppressed persons are at high risk; e.g., 1 - 5% of solid organ transplant recipients develop cryptococcosis (3). *C. neoformans* is found worldwide, while *C. gattii* is endemic to tropical and subtropical regions. In addition, hypervirulent strains of *C. gattii* have emerged in northwestern regions of North America, most notably on Vancouver Island (4–6).

Despite the need, there are no vaccines to protect humans from cryptococcosis (7). Populations which could be targeted for vaccination include persons: 1) HIV-infected; 2) on medications which suppress T cells (particularly transplant recipients); 3) living in *C. gattii*-endemic regions; and 4) with other high-risk diseases (e.g., sarcoidosis, lymphoma). Protection against experimental cryptococcosis can be obtained by immunization with cryptococcal strains missing virulence factors (such as chitosan, capsule, sterylglucosidase, and the F-box protein, Fbp1) or engineered to heterologously express interferon-γ (IFNγ) or overexpress the transcription factor Znf2 (8–15). In addition, protection can be obtained by immunizing mice with recombinant protein subunit vaccines adjuvanted in glucan particles (16, 17).

Previously, we deleted three chitin deacetylase (*CDA*) genes from the highly virulent KN99 strain of *C. neoformans*; the resulting strain, designated *cda1Δ2Δ3Δ*, is unable to produce chitosan, the deacetylated form of chitin, in the cryptococcal cell wall (11, 18). The *cda1Δ2Δ3Δ* strain is rapidly cleared from the lungs of CBA/J mice (11, 19). Mice that receive a pulmonary vaccination with live or heat-killed *cda1Δ2Δ3Δ* yeast cells are protected from a subsequent lethal lung challenge with the *C. neoformans* strain KN99 and (albeit to a lesser extent) *C. gattii* strain R265 (11, 20). Wild-type (WT) mice typically die within four weeks following infection with the virulent *C. neoformans* KN99 strain, with dissemination to the brain as is seen in most lethal human infections. In contrast, survival of vaccinated mice typically approaches 100%. In the present study, we explored the immunological basis for *cda1Δ2Δ3Δ* vaccine-mediated protection against pulmonary challenge with the *C. neoformans* KN99 strain.

## Results

### The kinetics of pulmonary clearance of live *cda1Δ2Δ3Δ* in CBA/J, BALB/c, C57BL/6, and NSG mice

We previously reported that the *cda1Δ2Δ3Δ* strain was rapidly cleared from the lungs of wild-type CBA/J mice (11, 19). In the first set of experiments, we extended the studies to determine the clearance rate of *cda1Δ2Δ3Δ* at a vaccination dose of 1×10^7^ CFU for CBA/J, BALB/c, C57BL/6, and <u>N</u>on-obese diabetic, <u>S</u>evere combined immunodeficiency, IL-2R common <u>G</u>amma-chain deficient (NSG) mice. The NSG mouse strain has particular translational relevance for use of live vaccines in immunocompromised populations, as it is important to demonstrate the vaccine strain is also avirulent in severely immunodeficient mice. NSG mice have multiple mutations making them highly immunodeficient. They lack mature B cells, T cells, and NK cells, and have defects in innate immunity and signaling pathways mediated by cytokine receptors which share the gamma chain (γ_c_) (21). When CBA/J, BALB/c, C57BL/6, and NSG mice were infected with 1×10^7^ *cda1Δ2Δ3Δ* yeast cells, CFUs in mouse lungs declined in a logarithmic manner to near undetectable levels over the 14d time course of the experiment (Figure 1).

**Figure 1.**
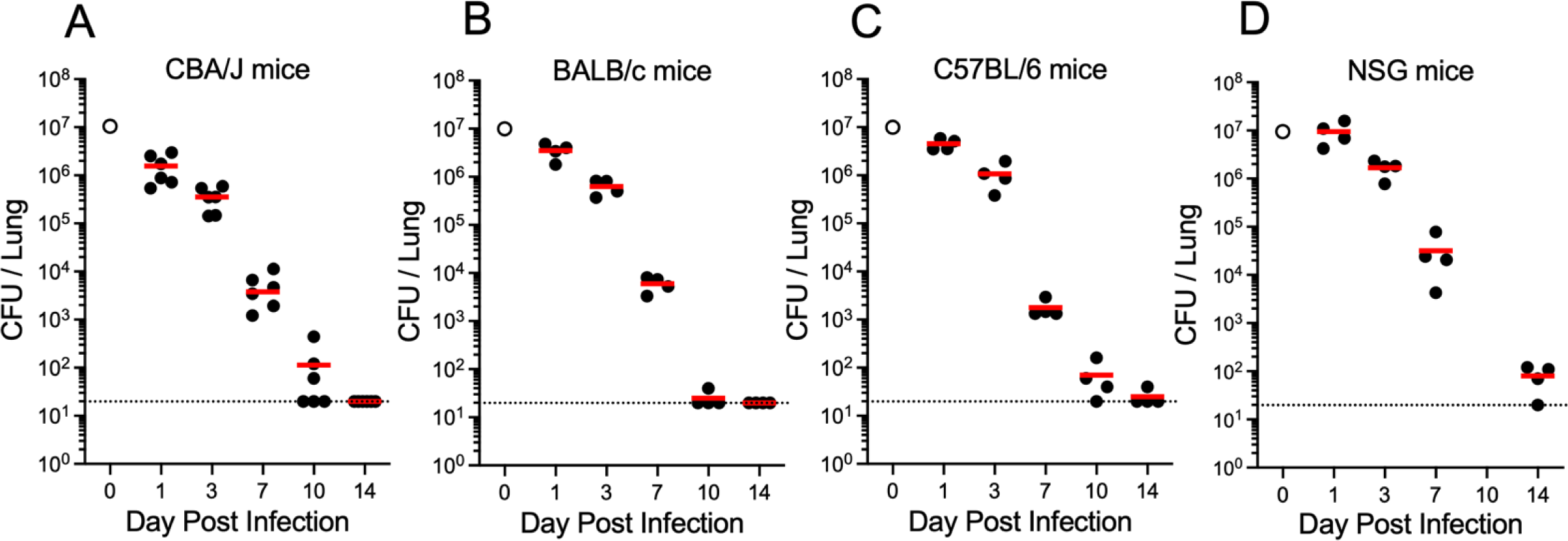
The kinetics of pulmonary clearance of cda1Δ2Δ3Δ in multiple mouse strains. CBA/J (A), BALB/c (B), C57BL/6 (C), and NSG (D) mice were infected OT with 1×10^7^ CFU of live cda1Δ2Δ3Δ strain. At designated days post infection, CFUs in the lungs were determined. Infected mice had n=4-6 mice per time point. Each circle represents CFUs in an individual mouse. Red horizonal bars denote mean values. Dotted lines at 20 CFU indicate the detection limit for CFU quantification. The inoculum is represented by open circle at day 0.

### Role of B cells and antibody in protection mediated by the *cda1Δ2Δ3Δ* vaccine

Having demonstrated that *cda1Δ2Δ3Δ* cells were avirulent in NSG mice that lacked B and T cells, we next systematically examined the contribution of B cells, CD8^+^ T cells, and CD4^+^ T cells to protection against cryptococcosis mediated by the live *cda1Δ2Δ3Δ* vaccine. The protective efficacy of the *cda1Δ2Δ3Δ* vaccine against pulmonary challenge with *C. neoformans* was examined in two strains of mice congenitally deficient in B cells. JHD mice (BALB/c background) lack mature B cells due to a targeted deletion of JH gene segments and are completely devoid of immunoglobulins (22). muMT mice (C57BL/6 background) lack B cells due to targeted disruption of the mu heavy chain gene (23); they make immunoglobulins at levels that are >100 fold lower than in normal mice (24).

Wild-type BALB/c and B cell-deficient JHD mice were vaccinated and then given a pulmonary challenge 42 days later with the virulent KN99 strain of *C. neoformans* after which they were monitored for survival (Fig 2A). Survival of both mouse strains was 100% when the experiment was terminated 70 days post infection (DPI) (Fig 2B). Moreover, when the experiment was terminated on day 70, lung CFUs were similar comparing WT and JHD mice (Fig 2C). Similar experiments were performed in wild-type C57BL/6 and muMT mice, except the mice received three vaccinations (orotracheal followed by two biweekly subcutaneous boosts) prior to infection (Fig 2D). The vaccine boosts were given because while a single vaccine dose confers significant protection to C57BL/6 mice, the protection is not as robust compared with other mouse strains (11). Wild-type C57BL/6 and muMT mice were protected by vaccination, although protection was less than 100% (Fig 2E). There was a trend towards decreased survival of the muMT mice compared to that seen in similarly vaccinated wild-type C57BL/6 mice. Lung CFUs in surviving mice did not significantly differ when comparing wild-type C57BL/6 and muMT mice. All unvaccinated mice, regardless of the mouse strain, succumbed within 30 DPI (Fig 2B, E).

**Figure 2.**
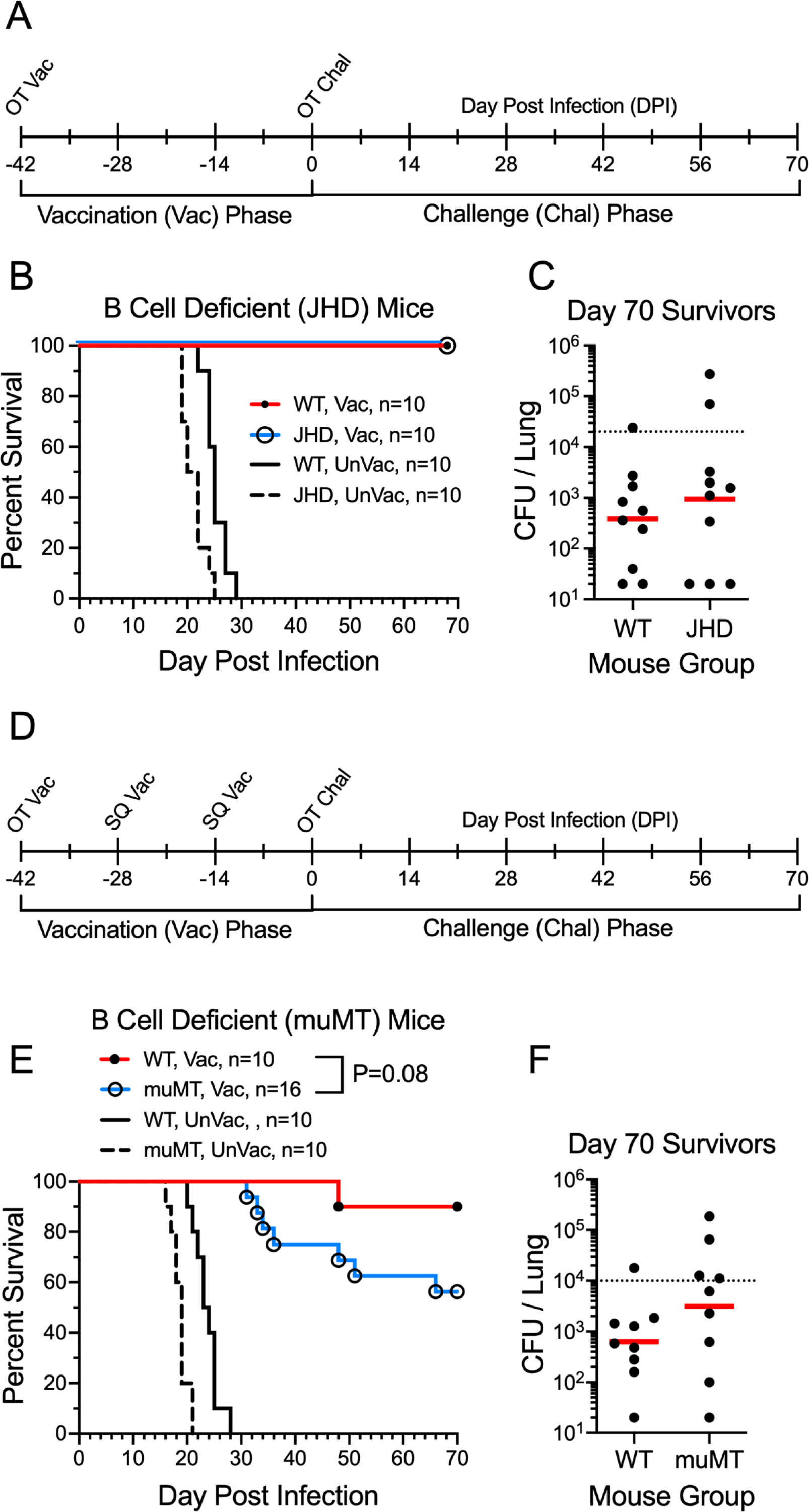
Contribution of B cells and antibody to protection following *cda1Δ2Δ3Δ* vaccination. (**A**) Protocol for experiments with wild-type (WT) BALB/c and JHD mice. Mice received orotracheal (OT) or intranasal (IN) vaccination with 1×10^7^ CFU live *cda1Δ2Δ3Δ* and challenge with 2×10^4^ CFU KN99. (**B**) Survival of vaccinated (Vac) and unvaccinated (UnVac) WT BALB/c and B cell deficient JHD mice were compared. (**C**) CFUs in the lungs of mice which survived to 70 DPI were determined. (**D**) Protocol for experiments with WT C57BL/6 and muMT mice. As in A except the mice also received two subcutaneous (SQ) boosts of 2×10^6^ CFU live *cda1Δ2Δ3Δ*. (**E**) Survival of vaccinated (Vac) and unvaccinated (UnVac) wild-type C57BL/6 and muMT mice were compared. **(F)** CFUs in the lungs of mice which survived to 70 DPI were determined. Kaplan-Meier survival curves were compared using the Mantel-Cox log rank test. For each strain of mice, p<0.001 comparing vaccinated with unvaccinated mice. Data are from ≥2 independent experiments. Number (n) of mice per group is indicated in the figure inset. For CFU, each circle represents the CFU of an individual mouse. Red horizontal bars denote mean values. Dotted lines indicate the KN99 challenge dose.

### Role of α/β T cells and CD8^+^ T cells in protection mediated by the *cda1Δ2Δ3Δ* vaccine

We next turned our attention to the role of T cells. TCRβ deficient mice (C57BL/6 background) contain a targeted deletion of T cell receptor β-chain and lack α/β T cells, but have normal differentiation of γ/δ T cells (25). The *cda1Δ2Δ3Δ* vaccine failed to protect TCRβ mice from a lethal challenge with *C. neoformans* (Fig 3A), strongly suggesting a critical role for α/β T cells in vaccine-mediated protection. As TCRβ mice lack both CD4^+^ and CD8^+^ α/β T cells, we next examined whether the CD8^+^ T cell subset contributed to protection by comparing survival following cryptococcal challenge of vaccinated wild-type C57BL/6 and β-microglobulin knock out (β2m) mice. β2m mice lack surface expression of major histocompatibility complex (MHC) class I and are virtually devoid of CD8^+^ T cells (26). The survival curves did not significantly differ; in fact, there was a non-significant trend towards increased survival in the β2m mice (Fig 3B).

**Figure 3.**
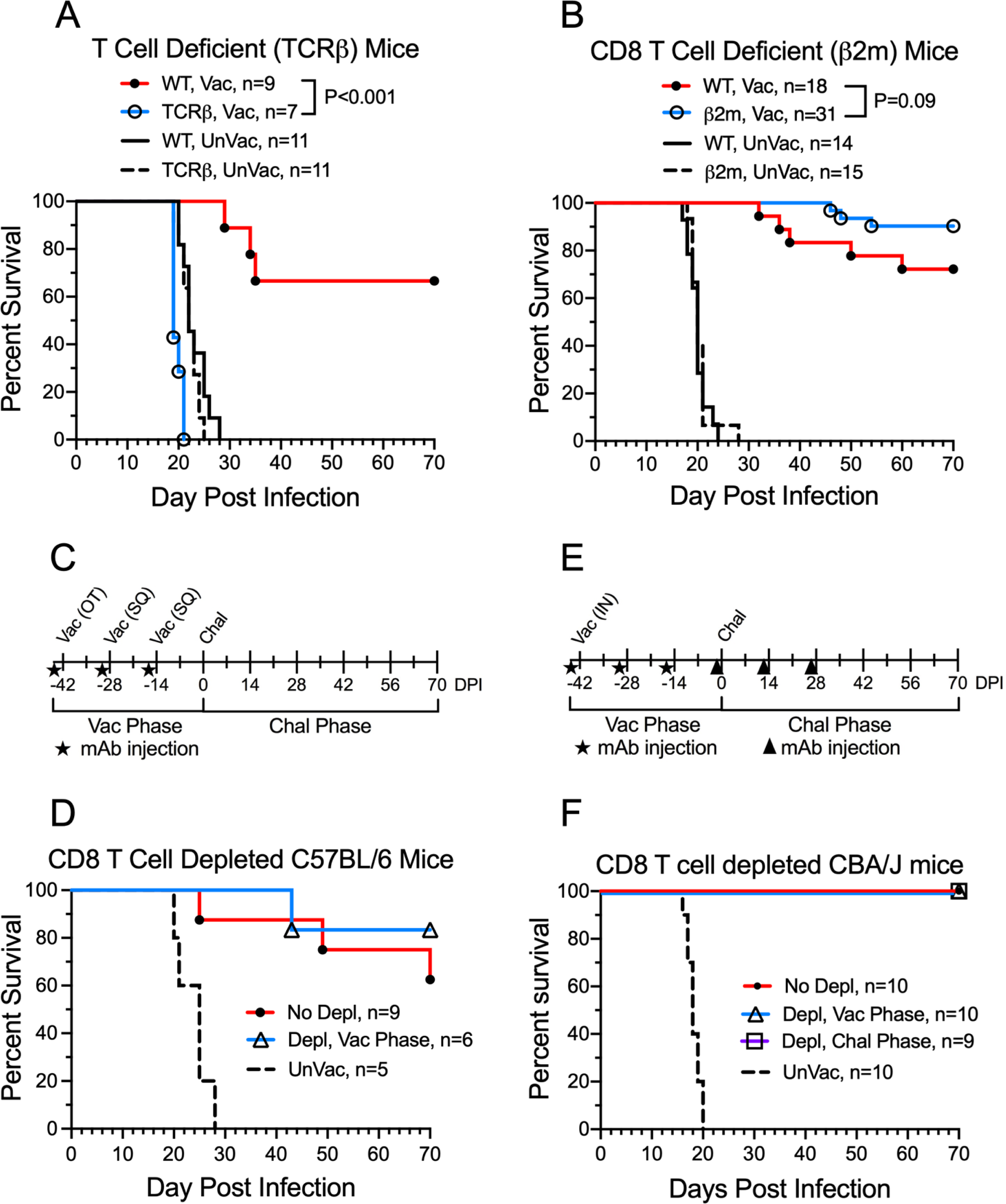
Contribution of CD8^+^ T cells to protection mediated by vaccination with *cda1Δ2Δ3Δ*. **(A)** Wild-type (WT) C57BL/6 mice and α/β T cell-deficient (TCRβ) mutant mice on the C57BL/6 background received an OT vaccination (1×10^7^ CFU) and two biweekly subcutaneous (SQ) boosts (2×10^6^ CFU each) with *cda1Δ2Δ3Δ*. Two weeks after the second SQ boost, mice were challenged with 1×10^4^ CFU KN99 and followed 70 days for survival. Vac = vaccinated with *cda1Δ2Δ3Δ*. UnVac = unvaccinated. **(B)** As in A except CD8^+^ T cell-deficient (B2m) mice were compared to WT mice. **(C**) Protocol for vaccination (Vac)/challenge (Chal) studies including administration of CD8^+^ T cell depleting mAb 2.43 mAbs by intraperitoneal injection. (**D,E**) Survival of vaccinated C57BL/6 (**D**) and CBA/J (**E**) mice following three bi-weekly injections of mAb 2.43 during the Vac phase or Chal phases. Statistics by Mantel-Cox log rank test. Data are from ≥2 independent experiments. Number (n) of mice per group is indicated in the figure inset.

β2m mice have perturbations aside from CD8 deficiency, including elevated iron levels, reduced neonatal Fc receptor function, and deficient natural killer T cell activity (27–29). As an alternative method to examine the role of CD8^+^ T cells, we treated mice with monoclonal antibodies (mAbs) 2.43 and YTS 169.4 to deplete CD8^+^ T cells from C57BL/6 mice and CBA/J mice, respecitvely (30). This also allowed us to expand studies on the importance of CD8^+^ T cells for protection to a second strain of mice, CBA/J. Confirmation that these mAbs were effective at depleting CD8^+^ T cells in C57BL/6 and CBA/J mice was shown by FACS (Supplementary Fig 1A and B). For the studies with C57BL/6 mice, mAb 2.43 was administered two days prior to each of the three vaccinations and then the mice were followed for survival following fungal challenge (Fig 3C and D). For the studies with CBA/J mice, three biweekly doses of mAb YTS 169.4 were administered either during the vaccine phase or during the challenge phase of the experiment (Fig 3E and F). Regardless of when CD8^+^ T cell depletion was performed, vaccinated C57BL/6 and CBA/J mice had >80% survival following *C. neoformans* challenge (Fig 3D and 3F). Moreover, survival was not significantly different comparing mAb-treated mice and mice with intact CD8^+^ T cell populations.

### Role of CD4^+^ T cells in protection mediated by the *cda1Δ2Δ3Δ* vaccine

The lack of a phenotype with mice lacking CD8^+^ T cells suggested CD4^+^ T cells were critical for *cda1Δ2Δ3Δ* vaccine-mediated protection. Therefore, we next examined vaccine-mediated protection in mice deficient in MHC class II (MHCII); these mice lack CD4^+^ T cells. MHCII mice vaccinated with *cda1Δ2Δ3Δ* were not protected against lethal challenge with *C. neoformans*; the survival curves of vaccinated and unvaccinated MHCII mice were practically superimposable (Fig 4A). To validate the observations about the importance of CD4^+^ T cells and to extend these observations to other mouse strains, we used the anti-CD4 mAb GK1.5 to deplete CD4^+^ T cells. Published data by our group and others demonstrated GK1.5 depletes the blood CD4^+^ T cell population within 24h of injection (31, 32). Moreover, depletion is sustained for at least two weeks, followed by a slow rebound in numbers (and, Supplementary Fig 1C and 1D). Three mouse strains, C57BL/6 (Fig 4B), CBA/J (Fig 4C), and BALB/c (Fig 4D) mice were studied. The experimental scheme was similar to what was used to deplete CD8^+^ T cells in the experiments depicted in Figure 3, except CBA/J and BALB/c mice received a intranasal or OT vaccination, but no subsequent SQ vaccinations. Mice received three injections of GK1.5 at two-week intervals during either the vaccination or challenge phase of the experiment. For each of the mouse strains, protection was lost when GK1.5 was administered during the vaccination phase. Importantly though, significant protection was retained if CD4^+^ T cells were depleted in the challenge phase. In BALB/c the protection was similar to the no depletion control, while in CBA/J and C57BL/6, the protection was lessened, but still significantly different than depletion in the vaccination phase. These data indicate that CD4^+^ T cells are essential for protection prior to vaccination, but are not absolutely required to sustain the protection.

**Figure 4.**
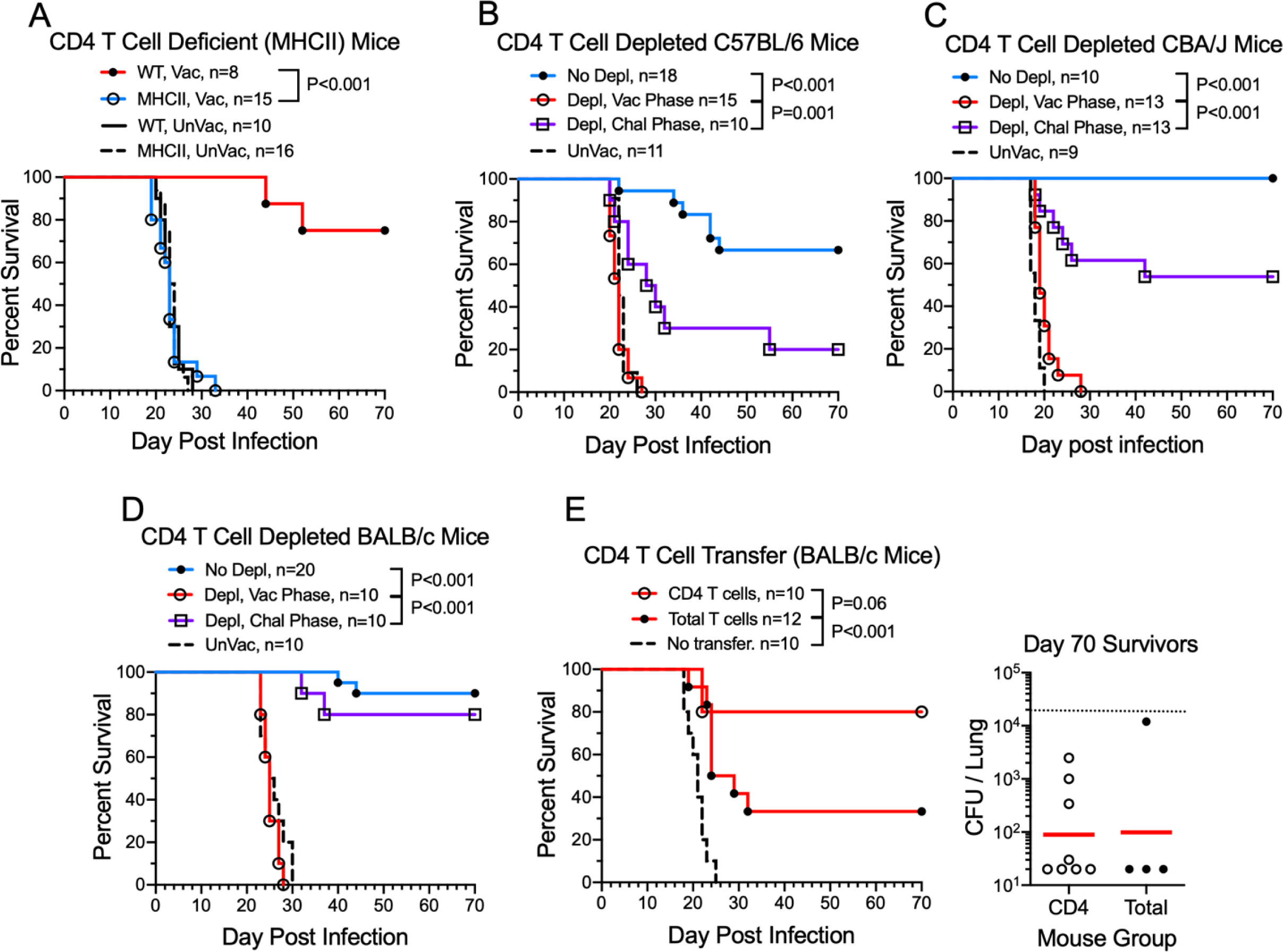
Contribution of CD4^+^ T cells to protection afforded by *cda1Δ2Δ3Δ* vaccination. (**A**) Vaccinated wild-type C57BL/6 (WT) and CD4^+^ T cell-deficient (MHCII) mice received three biweekly vaccinations (OT followed by two SQ boosts) with *cda1Δ2Δ3Δ*. Two weeks after the last boost, mice were challenged with 1×10^4^ CFU KN99. (**B,C,D**) Survival of vaccinated C57BL/6 (**B**), CBA/J (**C**), and BALB/c (**D**) mice following three bi-weekly injections of CD4^+^ T cell depleting mAb GK1.5 during the vaccinated (Depl, Vac) phase or challenged (Depl, Chal) phass. (**E**) Intravenous transfer of total T cells or CD4^+^ T cells purified from spleens of BALB/c mice vaccinated with *cda1Δ2Δ3* to naïve BALB/c mice followed by challenge with KN99. At 70 DPI, surviving mice were euthanized and lung CFUs of each mouse were determined. UnVac, unvaccinated. For CFU, each circle represents the CFU of an individual mouse. Red horizontal bars denote median values. The dotted line at 2×10^4^ indicates the KN99 challenge dose. Statistics by Mantel-Cox log rank test. Data are from ≥2 independent experiments. Number (n) of mice per group is indicated in the figure inset.

Finally, we performed adoptive transfer experiments to further assess the role of CD4^+^ T cells in vaccine-mediated protection (Fig 4E). Splenocytes were prepared from BALB/c mice vaccinated with the *cda1Δ2Δ3Δ* vaccine. Two groups of cells were purified, one consisting of total (CD3^+^) T cells and the other of the CD4^+^ T cell subpopulation. Adoptive intravenous transfer of each group of cells protected naïve mice against *C. neoformans* challenge, with the protection afforded by the purified CD4^+^ T cell population being more robust. Surviving mice showed similar clearance of KN99 from the lungs.

### The contribution of selected cytokines to *cda1Δ2Δ3Δ* vaccine-mediated protection

The finding that CD4^+^ T cells were critical to protection prompted us to explore the role of cytokines instrumental to their biasing and function. We focused on IFNγ, TNFα, and IL-23 as these three cytokines have known benefit in murine models of cryptococcosis (32–36). Vaccine-mediated protection was completely, or nearly completely, abrogated in mice lacking IFNγ (Fig 5A), IFNγR (Fig 5B), TNFα (Fig 5C), and IL-23p19 (Fig 5D).

**Figure 5.**
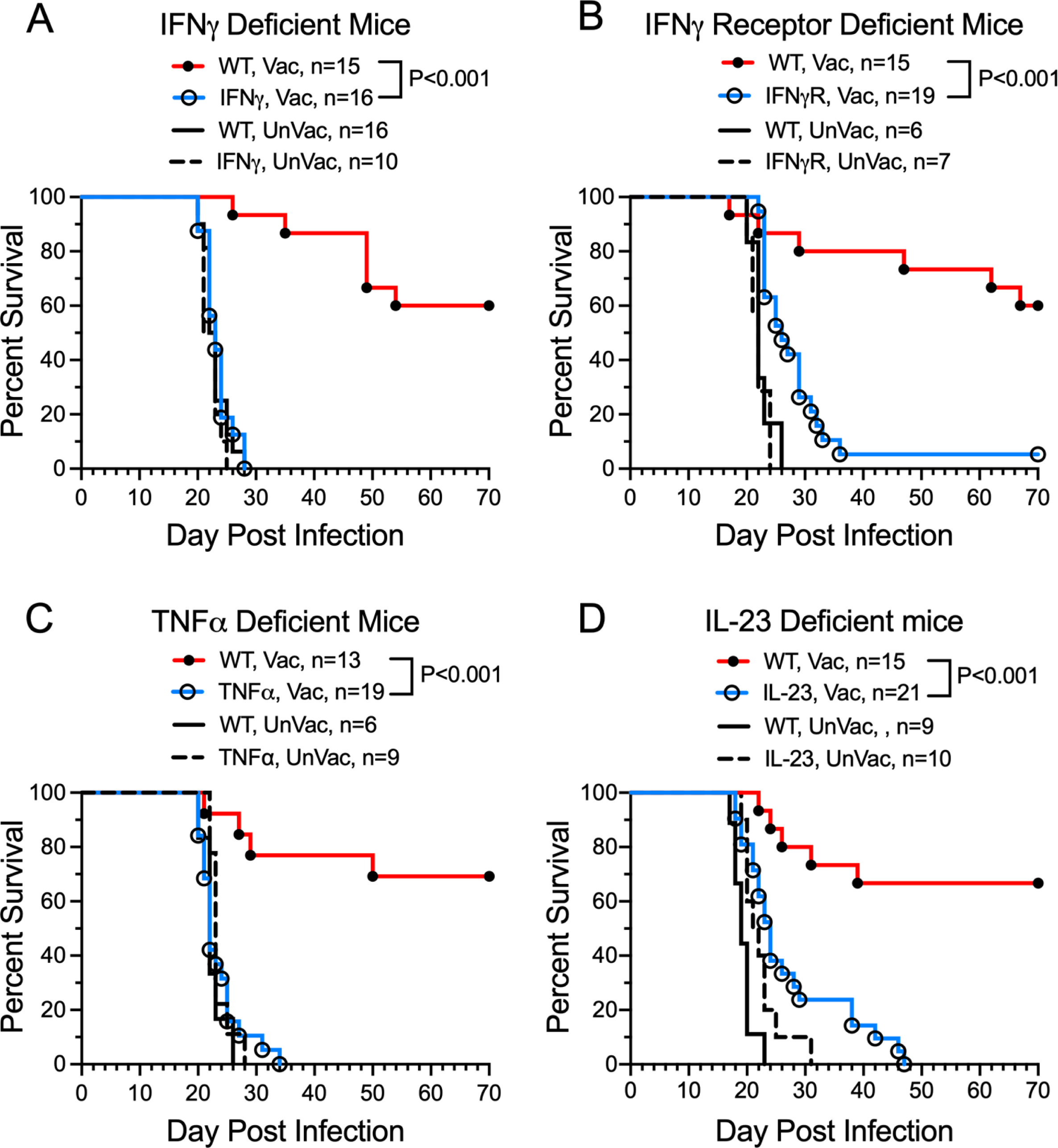
Contribution of selected cytokines to cda1Δ2Δ3Δ vaccine-mediated protection. Survival of vaccinated (Vac) and unvaccinated (UnVac) wild-type (WT) C57BL/6 mice were compared with that of mice deficient in (**A**) IFNγ, (**B**) IFNγR, (**C**) TNFα, and (**D**) IL-23p19 following an OT challenge with KN99. Statistics by Mantel-Cox log rank test. Data are from ≥2 independent experiments. Number (n) of mice per group is indicated in the figure inset.

### The nature of the lung T cell response following *cda1Δ2Δ3Δ* vaccination and infection

The above experiments established that CD4^+^ T cells and the cytokines IFNγ, TNFα, and IL-23 were required for vaccine-mediated infection. We next examined the quantity and quality of the pulmonary immune response following vaccination and/or infection. BALB/c mice were vaccinated with an OT dose of live *cda1Δ2Δ3Δ* vaccine and challenged six weeks later with *C. neoformans* KN99. Control mice were left unvaccinated and/or uninfected. Mice were euthanized either before challenge (0 DPI), or post challenge (10 DPI or 70 DPI) at which time their blood was collected by cardiac puncture, following which their lungs were harvested. As unvaccinated mice all die by 30 DPI, the 70 DPI group was comprised of only vaccinated mice.

Compared to unvaccinated mice at 10 DPI, vaccinated mice had significantly fewer CFUs in the lungs at 10 and 70 DPI (Fig 6A). However, total numbers of lung leukocytes, CD4^+^ T cells, and CD8^+^ T cells were not significantly different comparing unvaccinated and vaccinated mice at the 0 DPI and 10 DPI time points (Fig 6B-D). Notably, numbers of total leukocytes and CD4^+^ T cells increased in infected mice at 10 DPI and then returned to near baseline levels at 70 DPI. In contrast, CD8^+^ T cell counts did not significantly differ regardless of vaccination or infection status.

**Figure 6.**
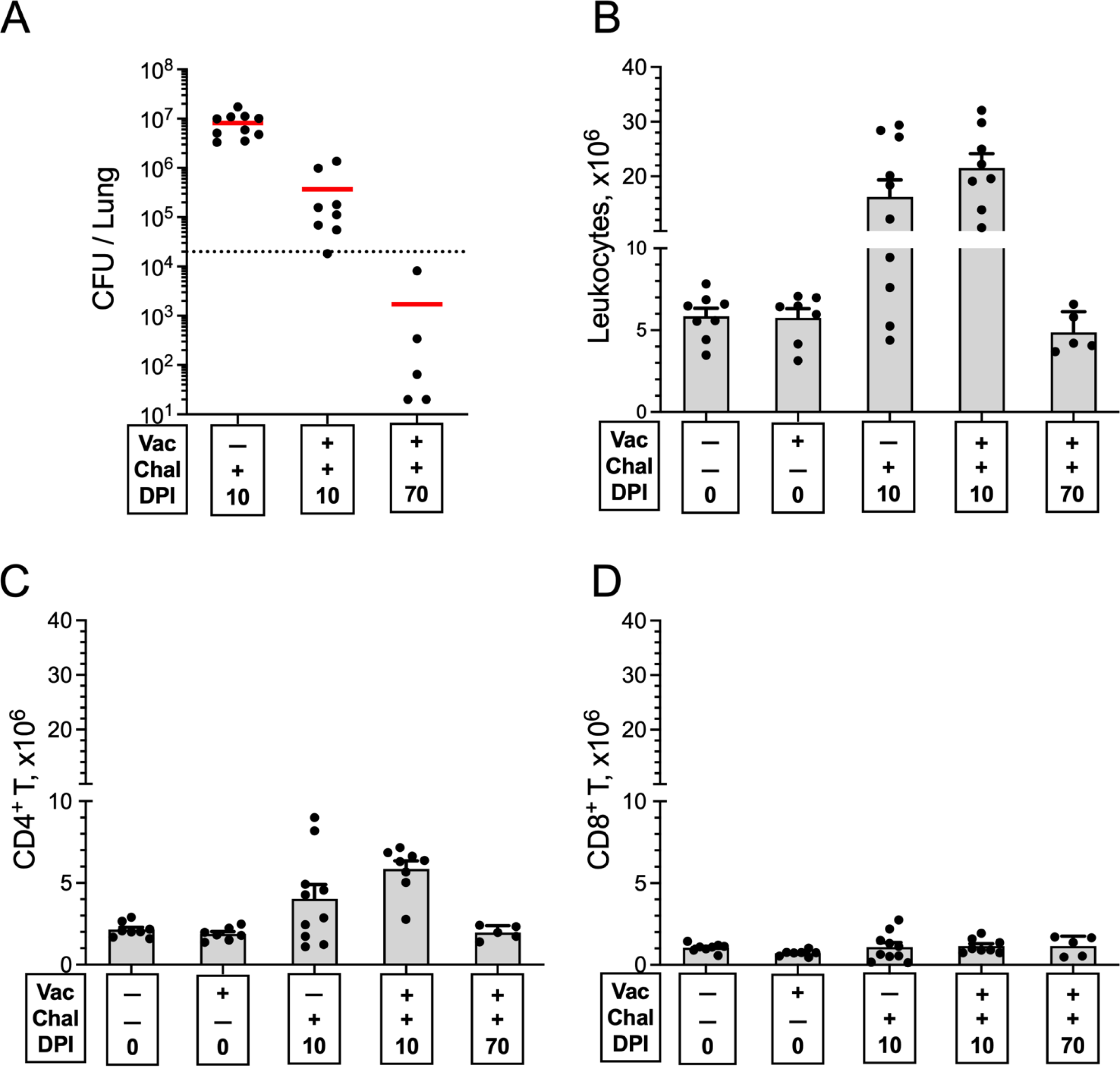
Quantitation of lung CFUs, leukocytes, and CD4+ and CD8+ T cells following vaccination and/or infection. BALB/c mice were vaccinated orotracheally with live *cda1Δ2Δ3Δ*. Six weeks later, the mice received a pulmonary challenge with KN99. Mice were euthanized at 0 DPI (uninfected), 10 DPI, or 70 DPI. Controls included unvaccinated mice euthanized at 0 DPI or 10 DPI. Lungs were harvested and single-cell suspensions were prepared. (A) CFUs per lung were determined. The horizontal bar represents the median CFUs. The dotted line depicts the challenge inoculum. (B) Leukocytes were purified on a Percoll gradient and counted. (C, D) The numbers of CD4^+^ and CD8^+^ T cells were calculated by multiplying the percentage of each population, as determined by FACS, times the total leukocyte count. Data are from two independent experiments, each with 4-6 mice/group (except for the 70 DPI group which had 2-3 mice/group). the data are presented as mean ± SEM. Vac, vaccinated with *cda1Δ2Δ3Δ*; Chal, challenged with *C. neoformans* KN99; DPI days post infection. Statistical comparisons between groups are shown in Supplementary Table 1.

Next, we measured expression of the activation marker CD154 and the intracellular cytokines IFNγ, TNFα, and IL-17A by lung CD4^+^ T cells following ex vivo stimulation with HK *cda1Δ2Δ3Δ* and HK KN99 (Fig 7). These three intracellular cytokines were chosen for study due to their importance in host defenses against cryptococcosis (32–35) and the loss of protection phenotype seen in vaccinated mice deficient in IFNγ, IFNγR, TNFα, or IL-23p19 (Fig 5). IL-23 is a key cytokine that promotes IL-17 production (37). Cells left unstimulated and stimulated with the superantigen SEB served as negative and positive controls, respectively.

**Figure 7.**
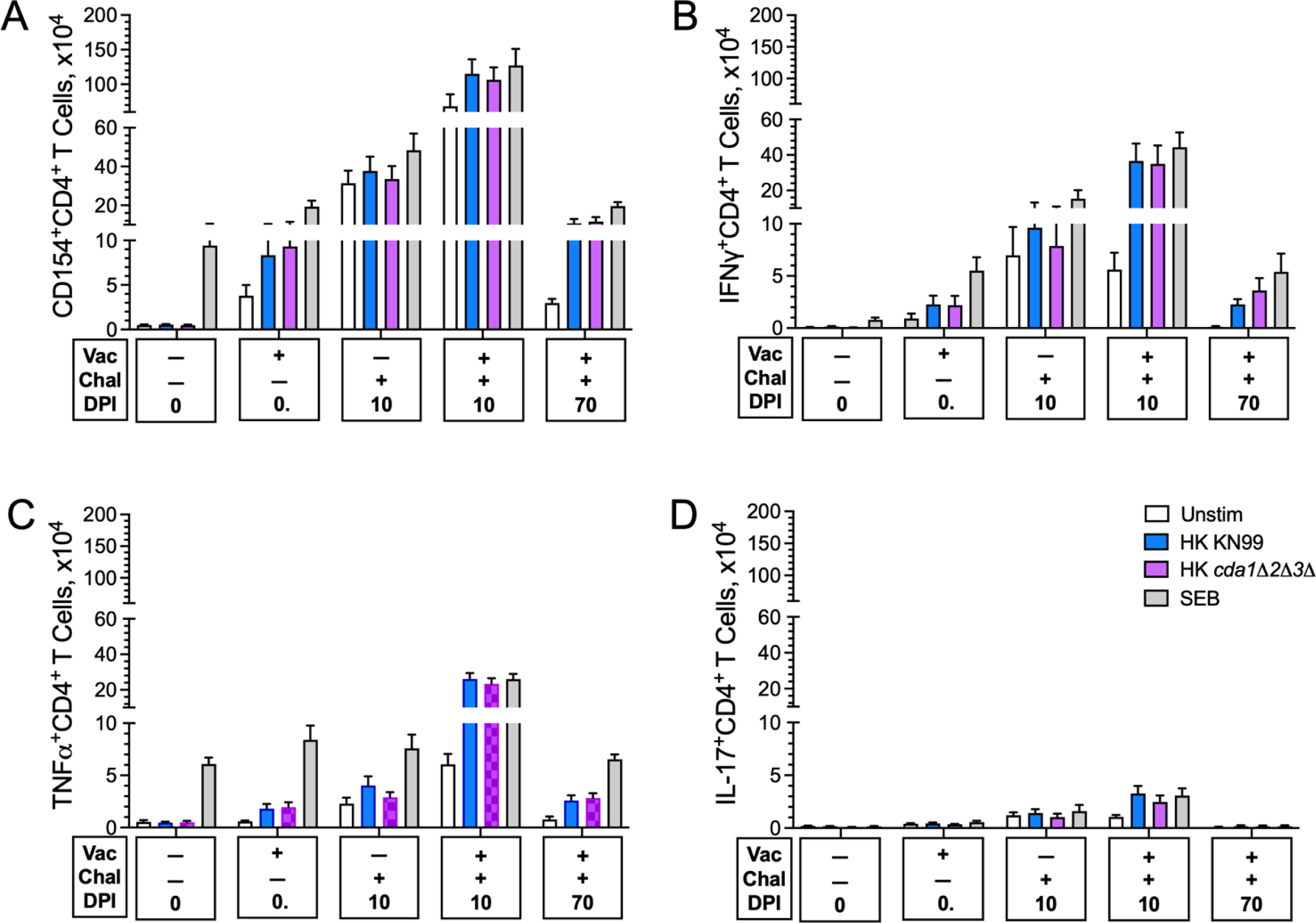
Ex vivo antigen-stimulated CD4^+^ T cell activation and intracellular cytokine production following vaccination and/or infection. Lung leukocytes from the experiment shown in Fig 6 were left unstimulated (Unstim) or stimulated with Staphylococcus enterotoxin B (SEB), HK KN99, or HK *cda1Δ2Δ3Δ* for 18h in complete media supplemented with 0.5 µg/ml amphotericin B. Brefeldin A was added during the last 4h of culture. The cells were then stained and analyzed by polychromatic FACS. The numbers of CD4^+^ T cells expressing the activation marker CD154 (A) or producing the intracellular cytokines IFNγ (B), TNFα (C) and IL17A (D) following ex vivo stimulation are shown. Data are from two independent experiments, each with 4-6 mice/group. Data are presented as mean ± SEM. Vac, vaccinated with *cda1Δ2Δ3Δ*; Chal, challenged with *C. neoformans* KN99; DPI, days post infection. Statistical comparisons between groups are shown in Supplementary Table 2.

In mice that were vaccinated but not infected, both *cda1Δ2Δ3Δ* and KN99 stimulated significant expression of CD154 (Fig 7A), IFNγ (Fig 7B), and TNFα Fig 7C) (but not IL-17A, Fig 7D) in the CD4^+^ T cell lung population. By 10 DPI, all four of these subpopulations of CD4^+^ T cells expanded, with more robust expansion observed in mice that were both vaccinated and infected compared with infected alone. Contraction of CD154, IFNγ, TNFα, and IL-17A expressing CD4^+^ T cells was observed in vaccinated mice that survived 70 DPI. For all groups, the numbers of CD4^+^ T cells expressing these four markers were similar following stimulation with HK KN99 compared with HK *cda1Δ2Δ3Δ*. Notably, modest increases over baseline numbers of CD154, IFNγ, TNFα, and IL-17A expressing CD4^+^ T cells were seen in groups that were vaccinated and/or infected but left unstimulated ex vivo. We postulate this is due, at least in part, to T cell activation and stimulation from residual fungal antigens in the lungs.

### IFNγ production by ex vivo stimulated lung leukocytes following vaccination and/or infection

Many cell types in addition to CD4^+^ T cells make IFNγ (38). To get a broader picture of pulmonary IFNγ production in vaccinated and infected mouse lungs, we examined IFNγ production in the supernatants of lung cells from vaccinated and/or infected mice stimulated 18h ex vivo with HK *cda1Δ2Δ3Δ* and HK KN99 (Fig 8A). As in Figure 7, unstimulated and SEB-stimulated cells served as controls. Undetectable to low levels of IFNγ were found in supernatants from unstimulated lung cells. Neither HK *C. neoformans* preparation stimulated appreciable IFNγ in unvaccinated mice. In contrast, HK *cda1Δ2Δ3Δ* and HK KN99 stimulated a vigorous IFNγ response in lung cells from vaccinated mice. Levels were even higher in mice that were vaccinated and infected. The high IFNγ levels persisted even at 70 DPI.

**Figure 8.**
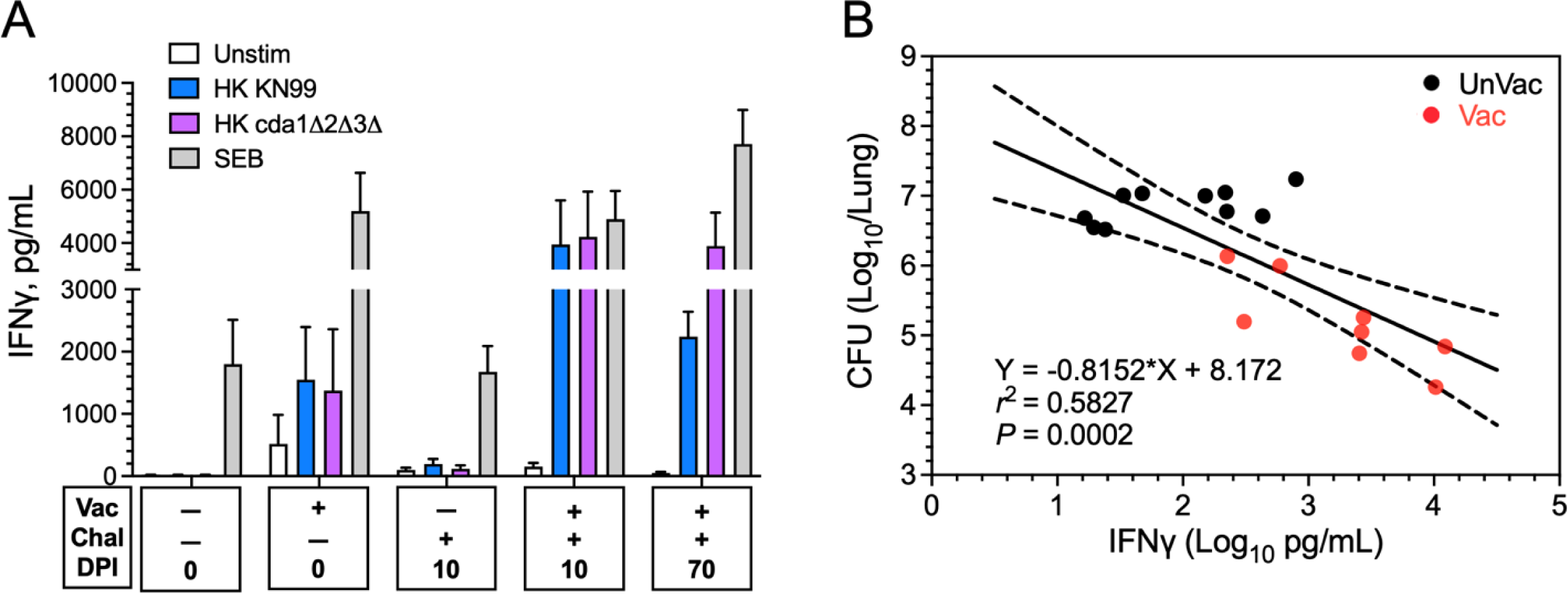
IFNγ production by ex vivo stimulated lung leukocytes following vaccination and/or infection. (A) Lung leukocytes were purified from vaccinated and/or infected mice and stimulated ex vivo for 18h using the same mice and protocol as in Fig 7. Supernatants were collected and analyzed for IFNγ by ELISA. Data are means ± SEM. Statistical comparisons between groups are shown in Supplementary Table 3. (B) Correlation between lung CFUs (see Fig 6A) and lung leukocytes IFNγ levels following HK KN99 stimulation in individual mice at 10 DPI. Data were analyzed with simple linear regression and are presented with best-fit line and confidence bands. Pearson correlation was used for statistical analysis. UnVac, unvaccinated; Vac vaccinated with live *cda1Δ2Δ3Δ*; Chal challenge with *C. neoformans* KN99; DPI, days post infection. Data are from two independent experiments, each with 4-6 mice/group.

To examine further the contribution of IFNγ to vaccine-mediated protection, we looked at the correlation of lung CFUs and IFNγ levels in vaccinated and unvaccinated mice at 10 DPI. A highly significant inverse correlation between CFUs and IFNγ levels was found (Fig 8B), lending further support for the critical contribution of IFNγ to vaccine-induced immunity.

## Discussion

In humans and experimental mouse models, CD4^+^ T cell immunity is required for protection against cryptococcosis (39). However, vaccines against cryptococcosis could work by eliciting protective responses from arm(s) of the immune system not in play during natural infection. Moreover, in situations where CD4^+^ T cells are deficient, immune plasticity could result in responses from cell types not required for protection in immunocompetent hosts. For example, an attenuated *Blastomyces dermatitidis* vaccine induced durable immunity in mice lacking CD4^+^ T cells by stimulating antigen-specific CD8^+^ T cell responses (40). Thus, it is important to determine immunological mechanisms of protection as that will be informative for identifying the patient populations for which the vaccine is most likely to be efficacious. Therefore, we systematically examined the components of the immune system which mediate protection to the *cda1Δ2Δ3Δ* vaccine.

A non-essential role for B cells was demonstrated as vaccine-mediated protection was retained in B cell-deficient muMT and JHD mice on the C57BL/6 and BALB/c backgrounds, respectively. These data suggest the *cda1Δ2Δ3Δ* vaccine does not require antibody for protection. However, an ancillary contribution of antibody to protection could not be ruled out as there was a trend towards increased mortality in the vaccinated muMT mice compared to their wild-type counterparts.

In vivo and in vitro roles for CD8^+^ T cells in host cryptococcal defenses have been demonstrated (41–43). However, our data demonstrate that CD8^+^ T cells were not required for protection mediated by the *cda1Δ2Δ3Δ* vaccine. First, the vaccine protected β2m^-/-^ mice, which are largely devoid of CD8^+^ T cells, from *C. neoformans* challenge as well as it did their wild-type counterparts. Second, mAb-mediated depletion of CD8^+^ T cells did not impact the survival of two strains of vaccinated mice.

In contrast, consistent with their paramount role in host defenses against cryptococcosis (3, 44), protection endowed by the *cda1Δ2Δ3Δ* vaccine was completely lost in mice congenitally deficient in CD4^+^ T cells. Moreover, adoptive transfer of unfractionated T cells or the CD4^+^ T cell subset from vaccinated mice conferred protection upon naïve mice. Depletion of CD4^+^ T cells using an anti-CD4 mAb abolished vaccine-mediated protection in mice if the depletion was performed at the time of vaccination. Importantly though, if CD4^+^ T cells were depleted after vaccination but prior to infection, then significant protection was retained. These results were consistent across three genetically distinct mouse strains; CBA/J, BALB/c, and C57BL/6, which have been described as highly resistant, moderately resistant, and susceptible, respectively, in mouse models of cryptococcosis (45). Thus, while CD4^+^ T cells are necessary to initiate a protective immune response, once the mice are immunized, effector cells are able to act in an apparently CD4^+^ T cell-independent manner to protect against fungal challenge. This plasticity has translational implications for persons with HIV/AIDS as it suggests vaccinating persons living with HIV while their CD4^+^ T cell counts are relatively high could provide protection should their CD4^+^ T cell counts then fall to the low levels (<100 cells/µL) associated with cryptococcosis (44). Studies in sub-Saharan Africa have found that the majority of patients presenting with AIDS-associated cryptococcosis had a prior history of being engaged in HIV care including receipt of antiretroviral drugs (46, 47). Similarly, *cda1Δ2Δ3Δ* vaccination could be offered to patients on transplant waiting lists in anticipation of their future need for immunosuppressive antirejection drugs (3). Protection against experimental cryptococcosis in the absence of CD4^+^ T cells has been described for other whole organism cryptococcal vaccines (15, 48). A caveat to the CD4^+^ T cell depletion experiments is resident memory lung T cells have been reported to be relatively resistant to depletion by anti-GK1.5 antibody (49).

Numerous murine and human studies have revealed IFNγ and TNFα are essential for host defenses against cryptococcosis (34, 39, 50–52). Indeed, the non-redundant contribution of these two cytokines in *cda1Δ2Δ3Δ*-mediated protection was clearly demonstrated as we found a complete loss of protection in mice null for IFNγ, IFNγR or TNFα. Our ex vivo studies identified Th1 cells as producers of these cytokines following *cda1Δ2Δ3Δ* vaccination and *C. neoformans* infection. It has been postulated that polyfunctional T cells which make IFNγ and TNFα have enhanced effector function and correlate with protection following vaccination (53). The data in BALB/c mice are consistent with our previous studies in CBA/J mice demonstrating that following infection with *C. neoformans*, *cda1Δ2Δ3Δ*-vaccinated mice had increases in pulmonary cytokines and chemokines often associated with Th1-type responses, including IFNγ and TNFα (11). Moreover, in the present studies, a significant inverse correlation was found between IFNγ levels and CFU in the lungs.

The heterodimeric cytokine IL-23 promotes differentiation and proliferation of IL-17-producing CD4^+^ cells (also known as Th17 cells) (37). Phenotypically, IL-23p19-deficient mice resemble mice deficient in IL-17 (54) although IL-23-independent production of IL-17 has been described (55). Protection mediated by the *cda1Δ2Δ3Δ* vaccine was almost completely lost in mice genetically deficient in the IL-23p19 subunit, suggesting IL-17 is integral to vaccine efficacy. Interestingly, the Th1 response was considerably more robust compared with the Th17 response following ex vivo antigenic stimulation of vaccinated and/or challenged mice. This suggests that cell types other than CD4^+^ T cells are making IL-17 or that IL-23 has protective effects independent of IL-17 in our vaccine model. Multiple cell types are known to produce IFNγ, TNFα, and IL-17a (56–58), and the contribution of individual cell types to vaccine immunity merits further study. Recently, IFNγ and IL-17 production by γδ T cells was shown to contribute to protection afforded by the *C. neoformans* Δsgl1 vaccine (59). In addition to IL-23, the p19 subunit reportedly associates with CD5 antigen-like (CD5L) protein; this p19/CD5L heterodimeric cytokine reportedly enhances the differentiation of GM-CSF-secreting CD4^+^ T cells (60). Future studies will need to parse the role of p19 and to examine whether vaccine protection is lost in mice specifically deficient in IL-17 or GM-CSF.

In summary, our data shed light on the complex immunological mechanisms required for protection against cryptococcosis afforded by the *cda1Δ2Δ3Δ* vaccine in mice. Several lines of evidence, when taken together, strongly suggest CD4^+^ T cells, particularly Th1 cells play a central role. First, protection was lost in mice deficient in CD4^+^ T cells, either due to a genetic mutation or mAb-mediated depletion. Second, protection could be restored by adoptive transfer of CD4^+^ T cells from vaccinated mice. Third, protection was lost in mice deficient in either IFNγ or IFNγR. Finally, mice vaccinated with *cda1Δ2Δ3Δ* and then infected with *C. neoformans* had robust recruitment of Th1 cells to the lungs. Non-redundant roles for TNFα and IL-23p19 were also seen. Importantly though, once vaccinated, mice were protected from an otherwise lethal challenge even if their CD4^+^ T cells were depleted. These results suggest a translational path for the *cda1Δ2Δ3Δ* vaccine; populations at high risk for cryptococcosis could be vaccinated while their CD4^+^ T cell function is relatively intact.

## Materials and Methods

### Reagents

Thermo Fisher Scientific (Pittsburgh, PA) was the source for reagents, except where noted. The buffer for flow cytometry staining was phosphate buffered saline (PBS) supplemented with bovine serum albumin (0.5%). Complete medium for cell culture was RPMI 1640 containing 10% fetal bovine serum (FBS), 10 mM HEPES, 2 mM L-alanyl-L-glutamine (GlutaMAX), 100 U/mL penicillin and 100 µg/mL streptomycin.

### Mouse strains

Table 1 lists the mouse strains used. CBA/J mice were purchased from Jackson Laboratories, and housed in the ABSL2 facilities at Washington University. Other mouse strains were bred and housed in a specific pathogen-free environment in the animal facilities at the University of Massachusetts Chan Medical School. Unless indicated, mice of both sexes were used in approximately equal numbers. All transgenic mouse strains were periodically genotyped and/or phenotyped. All animal procedures were carried out under protocols approved by the two schools’ Institutional Use and Care of Animals Committee.

**Table 1.** Mouse strains.

| **<u>Strain</u>** | **<u>Phenotype</u>** | **<u>Background</u>** | **<u>Source (reference)</u>** |
| --- | --- | --- | --- |
| C57BL/6 | Wild-type | C57BL/6 | JAX #000664 |
| BALB/c | Wild-type | BALB/c | JAX #000651 |
| CBA/J | Wild-type | CBA/J | JAX #000656 |
| NSG | Severely immunodeficient | NOD | JAX #005557 (21) |
| muMt | Mature B cell deficient | C57BL/6 | JAX #002288 (64) |
| Jh (JHD) | Mature B cell deficient | BALB/c | Taconic #1147 (22) |
| TCRβ | $\alpha/\beta$ T cell receptor deficient | C57BL/6 | JAX #002118 (25) |
| β2m | CD8 T cell deficient | C57BL/6 | JAX #002087 (26) |
| MHCII | CD4 T cell deficient | C57BL/6 | JAX #003239 (65) |
| IFNγ | IFNγ deficient | C57BL/6 | JAX #002286 (66) |
| IFNγR | IFNγ receptor 1 deficient | C57BL/6 | JAX #003288 (67) |
| TNFα | TNFα deficient | C57BL/6 | JAX #003008 (68) |
| IL-23p19 | IL-23p19 deficient | C57BL/6 | Genentech (54) |

JAX, The Jackson Laboratory. NSG, Non-obese diabetic, Severe combined immunodeficiency, IL-2R common Gamma-chain deficient. muMT, mu (IgM) immunoglobulin heavy chain. TCRβ , T cell receptor beta chain. β2m, $\beta$ 2 microglobulin. MHCII, major histocompatibility locus class II. Catalog numbers of the mouse strains are included with their sources and references, where relevant.

### Vaccinations and infections

Vaccinations and infections were with the avirulent *cda1Δ2Δ3Δ* strain (18) and the highly virulent KN99α strain of *C. neoformans* (61), respectively. Studies with wild-type and genetically modified mice C57/BL6 and BALB/c were performed at UMass Chan Medical School. Stocks of the strains were maintained at -80°C as a mix of a yeast extract-peptone-dextrose (YPD) liquid culture shaken for 2d at 30°C combined 1:1 (v/v) with 50% sterile glycerol. Prior to use in mouse studies each stock was first grown on YPD agar medium at 30°C for 2-3d and used as inoculum of liquid YPD cultures; plates were stored at 4°C for up to 30d.

For the first vaccination, *cda1Δ2Δ3Δ* was shaken (225 rpm) in 25 mL of YPD in a 250 mL polycarbonate flask (CELLTREAT Scientific Products, Pepperell, MA) for 2d at 30°C; for subsequent vaccinations 14 mL polypropylene round-bottom tubes (Corning) and 4 mL YPD medium were used. Cells were collected by centrifugation for 5 min at 425xg, washed once with PBS (equal in volume to the culture medium) and suspended in 10 mL (flask culture) or 4 mL (tube culture) of PBS. The suspended cells were then diluted 1:100 in PBS and counted using a TC20 automated cell counter (Bio-Rad Laboratories, Hercules CA). For the first vaccination with *cda1Δ2Δ3Δ* (given to CBA/J, BALB/c and C57BL/6 mice), the cell count was adjusted with PBS to 2×10^8^ cells/mL. For the second and third vaccinations (administered to C57BL/6 mice only), the cell count was adjusted to 2×10^7^ cells/mL. Mice were challenged with KN99 that had been cultured in 4 mL YPD medium and shaken for 18h at 30°C. Cells were collected and washed, as above; the final concentration of KN99 depended on the strain of mice: C57BL/6 mice was 2×10^5^ cells/mL, BALB/c was 4×10^5^ cells/mL, and CBA/J mice was 2×10^6^ cells/mL of PBS.

For the first vaccination with *cda1Δ2Δ3Δ*, and for infection with KN99, mice were individually anaesthetized with isoflurane, USP (UMASS Department of Animal Medicine). Then 50 µL of the 2×10^8^ cells/mL suspension, (described above) was administered by orotracheal (OT) inoculation into the lungs (62). The second and third vaccinations of *cda1Δ2Δ3Δ* were by subcutaneous (SQ) injection of 100 µL in the abdomen. KN99 was also introduced into the lungs by orotracheal inoculation of 50 µL.

The intranasal vaccinations and infections with CBA/J mice were performed at Washington University, as described (62, 63). Yeast cells (*cda1*Δ*2*Δ*3*Δ and KN99) were cultivated in YPD medium (50 mL of the medium in a 250 mL flask) and shaken at 300 rpm for 48h at 30°C. Cells were collected by centrifugation at 3000xg for 10 min, and the pellet was washed twice with endotoxin-free PBS. PBS was added to the pellet to result in a final cell concentration of 2×10^8^ cells/mL. To inoculate intranasally, appoximately 5–6-week-old female CBA/J mice were anesthetized with an intraperitoneal injection (200 μL) of ketamine (8 mg/mL)/dexdomitor (0.05 mg/mL) mixture and then intranasally infected with 50 μL of the yeast cell suspension. After 10 min, the mice were administered with the reversal agent atipamezole (200 μL; 0.25 mg/mL) intraperitoneally. Reagents used for anesthesia were provided by the Washington University Division of Comparative Medicine Pharmacy.

For all infection experiments, mice were monitored daily; those showing signs of morbidity (including weight below 80% of pre-inoculation weight or extension of the cerebral portion of the cranium) were euthanized by CO_2_ asphyxiation and cervical dislocation. Survival studies were terminated at 70 DPI.

### Cryptococcal colony forming units (CFUs)

CFUs were determined for *cda1Δ2Δ3Δ* and KN99 inocula administered to the lungs to corroborate their respective cell counts. Lung CFUs were measured following homogenization of lung lobes in 4 mL PBS containing 200 U/mL penicillin and 200 µg/mL streptomycin using an OMNI TH homogenizer with tip adaptor for 7 mm x 110 mm hard tissue probe (OMNI International, Kennesaw, GA). Dilutions were plated on Sabouraud dextrose agar (Remel) or YPD agar and plates were incubated for 2-3 days at 30°C prior to counting. The total CFU per organ was computed, with a detection limit of 20 CFU per organ.

CD4^+^ and CD8^+^ T cell depletions

T cells were depleted from mice using monoclonal antibodies (mAb) GK1.5 for CD4^+^ T cells (Bio X Cell, Lebanon, NH), and mAb 2.43 (Cell Culture Company, Minneapolis, MN) or YTS 169.4 (Bio X Cell) for CD8^+^ T cells. Each mAb was administered every two weeks by intraperitoneal (IP) injection of 200 µg in 100 µL PBS. Timelines for IP injections are depicted in Figure 3.

### Transfer of T cells from vaccinated mice to naïve mice

BALB/c mice were vaccinated three times with *cda1Δ2Δ3*Δ. The first vaccination was OT and the second and third vaccinations were SQ, as described above. Spleens and lymph nodes were collected from 3-5 mice euthanized four weeks after the 3^rd^ vaccination. Total T cells and CD4^+^ T cells were each purified by negative selection following the manufacturer’s instructions provided with the mouse Pan T Cell Isolation Kit and CD4^+^ T Cell Isolation Kit, respectively (Miltenyi Biotec, Bergisch Gladbach, Germany). Purified T cells were transferred to naïve BALB/c mice via the tail vein; each mouse was injected with 5×10^6^ cells in 100 µL PBS. Two days after transfer the mice were infected with 2×10^4^ KN99 cells.

### Ex vivo stimulation and analysis of lung leukocytes

Lungs were collected and lung populations stimulated and analyzed as described (32), with minor modifications as noted. Briefly, lungs were collected after exsanguination by cardiac puncture and rinsed with PBS. Single cell suspensions were prepared using the Miltenyi Lung Dissociation Kit (Miltenyi Biotec). Leukocytes were enriched and *C. neoformans* were depleted following centrifugation on a Percoll gradient (40%/67%, Cytiva, Uppsala, Sweden). Lung cells (4 x 10^5^ cells/well) were cultured for 18 hours in 96-well flat bottom plates containing 200 µL complete media supplemented with 0.5 µg/ml amphotericin B. Stimuli included *Staphylococcus* enterotoxin B (SEB, 1 µg/ ml, Toxin Technology Inc, Sarasota, FL), heat-killed (HK) KN99 (50 µg/ml), and HK *cda1Δ2Δ3Δ* (*Cda123*, 50 µg/mL) for 18h in complete media supplemented with amphotericin B (0.5 µg/mL). Brefeldin A (5 µg/mL, BioLegend, San Diego, CA) was added during the last 4h of culture.

Plates were centrifuged at 825xg for 5 min and supernatants collected for IFNγ ELISA (R&D Systems Mouse IFNγ DuoSet ELISA Kit, Minneapolis, MN). Samples (50 µL) were assayed following the manufacturer’s instructions. The lower limit of detection of IFNγ was 10 pg/mL; values below the lower limit were assigned concentrations of 9 pg/mL. The leukocytes in the cell pellet were suspended and stained with the LIVE/DEAD Green Fixable Dead Cell Stain Kit (Invitrogen), followed by CD3-PE, CD4-PerCP/Cyanine5.5, and CD8-APC antibodies (BioLegend). Cells were then fixed and permeabilized using the Intracellular Fixation & Permeabilization Buffer Set, FcRs were blocked with rat anti-mouse CD16/CD32 monoclonal antibody 2.4G2 (BD, Franklin Lakes, NJ), and then the cells were stained with antibodies CD154-PE/Cyanine7, IFNγ-BV650, IL-17A-BV510, and TNFα-APC/Cyanine7 (BioLegend). Antibody catalog numbers and their dilutions were the same as in Wang et al. (32). A 5-laser Bio-Rad ZE5 flow cytometer (Bio-Rad) was used to acquire data. Data was analyzed using FlowJo version 10.10 software (BD). Gating was established using fluorescence minus one (FMO) controls and isotype controls, as illustrated in Supplementary Figure 2.

### Statistics

GraphPad Prism Version 10.1.2 (GraphPad Software, La Jolla, CA) was used for statistical analyses and drawing graphs. The Mantel-Cox log-rank test was used to assess significance when comparing Kaplan-Meier survival curves. Lung CFUs from the ex vivo studies had a nonparametric distribution; groups were compared using the Mann-Whitney test. Lung leukocytes, CD4^+^ T cells, and CD8^+^ T cells were normally distributed; groups were compared using one-way ANOVA with Tukey’s correction for multiple comparisons. Quantification of cytokine-producing CD4^+^ T cell numbers or levels of IFNγ within supernatants post ex vivo stimulation were presented as mean ± SEM, and statistical comparisons were conducted using two-way ANOVA with either Tukey’s or Dunnett’s correction for multiple comparisons. For the correlation analysis between lung CFUs and IFNγ levels, the data underwent simple linear regression using Pearson correlation, and were presented with a best-fit line and lines depicting 95% confidence intervals. A P value of <0.05 following corrections for multiple comparisons was considered significant.

## Acknowledgments

The authors thank Brian Maybruck for verifying some of the CD4^+^ T cell depletions.

This work was supported by National Institute of Allergy and Infectious Diseases, National Institutes of Health grants R01AI125045 (to J.K.L., C.A.S., and S.M.L.), and AI172154 (to S.M.L.), and contract 75N93019C00064 (to S.M.L.), all from the (NIH). M.H. was partially supported by NIH Training Grant T32 AI095213.

